# A familial Alzheimer’s disease-like mutation in the zebrafish *presenilin 1* gene affects brain energy production

**DOI:** 10.1101/542134

**Authors:** Morgan Newman, Nhi Hin, Stephen Pederson, Michael Lardelli

**Author notes:** Equal first authors (Morgan Newman, Nhi Hin). **Address for all authors:** University of Adelaide School of Biological Sciences Department of Molecular and Biomedical Sciences North Terrace, Adelaide, SA 5005 Australia.

## Abstract

To prevent or ameliorate Alzheimer’s disease (AD) we must understand its molecular basis. AD develops over decades but detailed molecular analysis of AD brains is limited to postmortem tissue where the stresses initiating the disease may be obscured by compensatory responses and neurodegenerative processes. Rare, dominant mutations in a small number of genes, but particularly the gene *PRESENILIN 1* (*PSEN1*), drive early onset of familial AD (EOfAD). Numerous transgenic models of AD have been constructed in mouse and other organisms, but transcriptomic analysis of these models has raised serious doubts regarding their representation of the disease state. Since we lack clarity regarding the molecular mechanism(s) underlying AD, we posit that the most valid approach is to model the human EOfAD genetic state as closely as possible. Therefore, we sought to analyse brains from zebrafish heterozygous for a single, EOfAD-like mutation in their *PSEN1*-orthologous gene, *psen1*. We previously introduced an EOfAD-like mutation (Q96_K97del) into the endogenous *psen1* gene of zebrafish. Here, we analysed transcriptomes of young adult (6-month-old) entire brains from a family of heterozygous mutant and wild type sibling fish. Gene ontology (GO) analysis revealed effects on mitochondria, particularly ATP synthesis, and on ATP-dependent processes including vacuolar acidification.

## Background

AD is the most common form of dementia with severe personal, social, and economic impacts. Rare, familial forms of AD exist caused by autosomal dominant mutations in single genes (reviewed by (1)). The majority of these mutations occur in the gene *PRESENILIN 1* (*PSEN1*) that encodes a multipass integral membrane protein involved in intra-membrane cleavage of numerous proteins (1).

A wide variety of transgenic models of AD have been created and studied. These are aimed at reproducing histopathologies posited to be central to the disease process, i.e. amyloid plaques and neurofibrillary tangles of the protein MAPT (2). However, analysis of transcriptome data derived from a number of these mouse models shows little concordance with transcriptomic differences between human AD brains and age-matched controls (3). We posit that, in the absence of an understanding of the molecular mechanism(s) underlying AD, the most objective approach to modeling this disease (or, at least, modeling its genetic form, EOfAD) is to create a genetic state as similar as possible to the EOfAD state in humans. Mouse “knock-in” models of EOfAD mutations were created over a decade ago and showed subtle phenotypic effects but not the desired histopathologies (e.g. (4, 5)). However, at that time, researchers did not have access to contemporary ‘omics technologies and transcriptome analysis of these models was never performed.

In humans, AD is thought to develop over decades and the median survival to onset age for EOfAD mutations in human *PSEN1* considered collectively is 45 years (6). Functional MRI of human children carrying EOfAD mutations in *PSEN1* has revealed differences in brain activity compared to non-carriers in individuals as young as 9 years of age (7). Presumably therefore, heterozygosity for EOfAD mutations in *PSEN1* causes early molecular changes/stresses that eventually lead to AD.

Transcriptome analysis is currently the most detailed molecular phenotypic analysis possible on cells or tissues. Here we present an initial analysis of the transcriptomic differences caused in young adult (6-month-old) zebrafish brains by the presence of an EOfAD-like mutation in the gene *psen1* that is orthologous to the human *PSEN1* gene. GO analysis supports very significant effects on mitochondrial function, especially synthesis of ATP, and on ATP-dependent functions such as the acidification of lysosomes that are critical for autophagy.

## Materials and Methods

The mutant allele, Q96_K97del, of *psen1* was a byproduct identified during our introduction of the K97fs mutation into *psen1* (that models the K115fs mutation of human *PSEN2* – see (8) for an explanation).

Q96_K97del is a deletion of 6 nucleotides from the coding sequence of the *psen1* gene. This is predicted to distort the first lumenal loop of the Psen1 protein. In this sense, it is similar to a number of EOfAD mutations of human *PSEN1* (9). Also, in common with all the widely distributed EOfAD mutations in *PSEN1*, (and consistent with the PRESENILIN EOfAD mutation “reading frame preservation rule” (1)), the Q96_K97del allele is predicted to encode a transcript that includes the C-terminal sequences of the wild type protein. Therefore, as a model of an EOfAD mutation, it is superior to the K97fs mutation in *psen1* (8).

To generate a family of heterozygous Q96_K97del allele (i.e. *psen1*^*Q96_K97del*^/+) and wild type (+/+) sibling fish, we mated a *psen1*^*Q96_K97del*^/+ individual with a +/+ individual and raised the progeny from a single spawning event together in one tank. Zebrafish can live for up to 5 years but, in our laboratory, typically show greatly reduced fertility after 18 months. The fish become fertile after around three months of age, so we regard 6-month-old fish as equivalent to young adult humans. Therefore we analysed the transcriptomes of entire young adult, 6-month-old fish brains using poly-A enriched RNA-seq technology, and estimated gene expression from the resulting single-end 75bp reads using the reference GRCz11 zebrafish assembly transcriptome (10, 11). Each zebrafish brain has a mass of approximately 7 mg. Since AD is more prevalent in human females than males, and to further reduce gene expression “noise” in our analyses, we obtained brain transcriptome data from four female wild type fish and four female heterozygous mutant fish. This data has been made publicly available at the Gene Expression Omnibus (GEO, see under *Availability of data and material* below).

## Results

### Differentially expressed genes (DE genes)

Genes differentially expressed between wild type and heterozygous mutant sibling fish were identified using moderated *t*-tests and a false discovery rate (FDR)-adjusted *p*-value cutoff of 0.05 as previously described (8, 12, 13). In total, 251 genes were identified as differentially expressed (see Additional File 1). Of these, 105 genes showed increased expression in heterozygous mutant brains relative to wild type sibling brains while 146 genes showed decreased expression.

### GO analysis

To understand the significance for brain cellular function of the differential gene expression identified in young adult heterozygous mutant brains we used the *goana* function (14) of the *limma* package of Bioconductor software (13) to identify GOs in which the DE genes were enriched at an FDR-corrected p-value of less than 0.05. 78 GOs were identified of which 20 addressed cellular components (CC). Remarkably, most of these CCs concerned the mitochondrion, membranes, or ATPases. 17 GOs addressed molecular functions (MF) and largely involved membrane transporter activity, particularly ion transport and ATPase activity coupled to such transport (Table 1). 41 GOs addressed biological processes (BP) and involved ATP metabolism, ribonucleoside metabolism, and transmembrane transport processes including vacuolar acidification (that has previously been identified as affected by EOfAD mutations in *PSEN1* (15)). Overall, our GO analysis indicates that this EOfAD-like mutation of zebrafish *psen1* has very significant impacts on cellular energy metabolism and transmembrane transport processes.

**Table 1.**
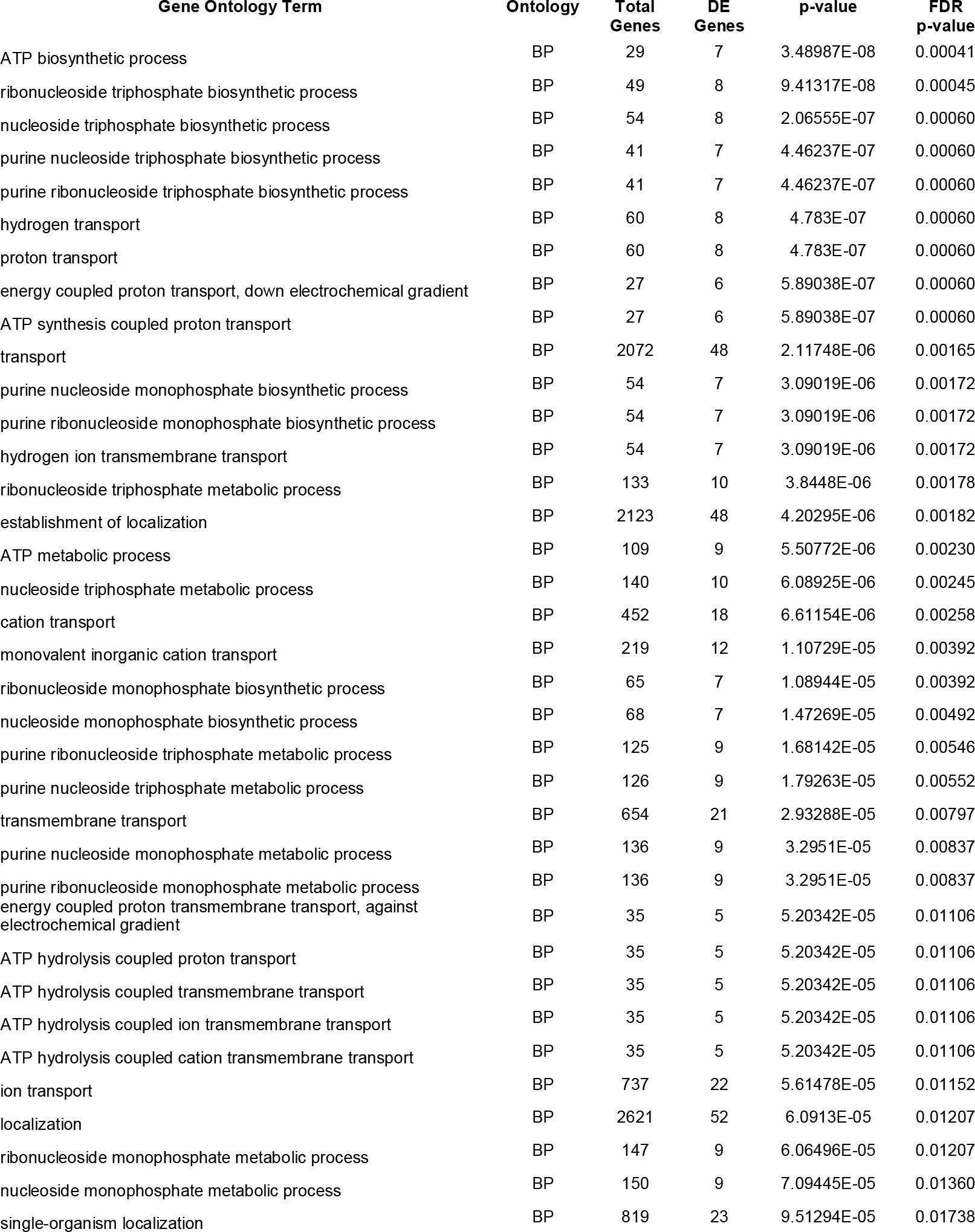
GOs enriched for genes differentially expressed between heterozygous mutant and wild type sibling fish brains.

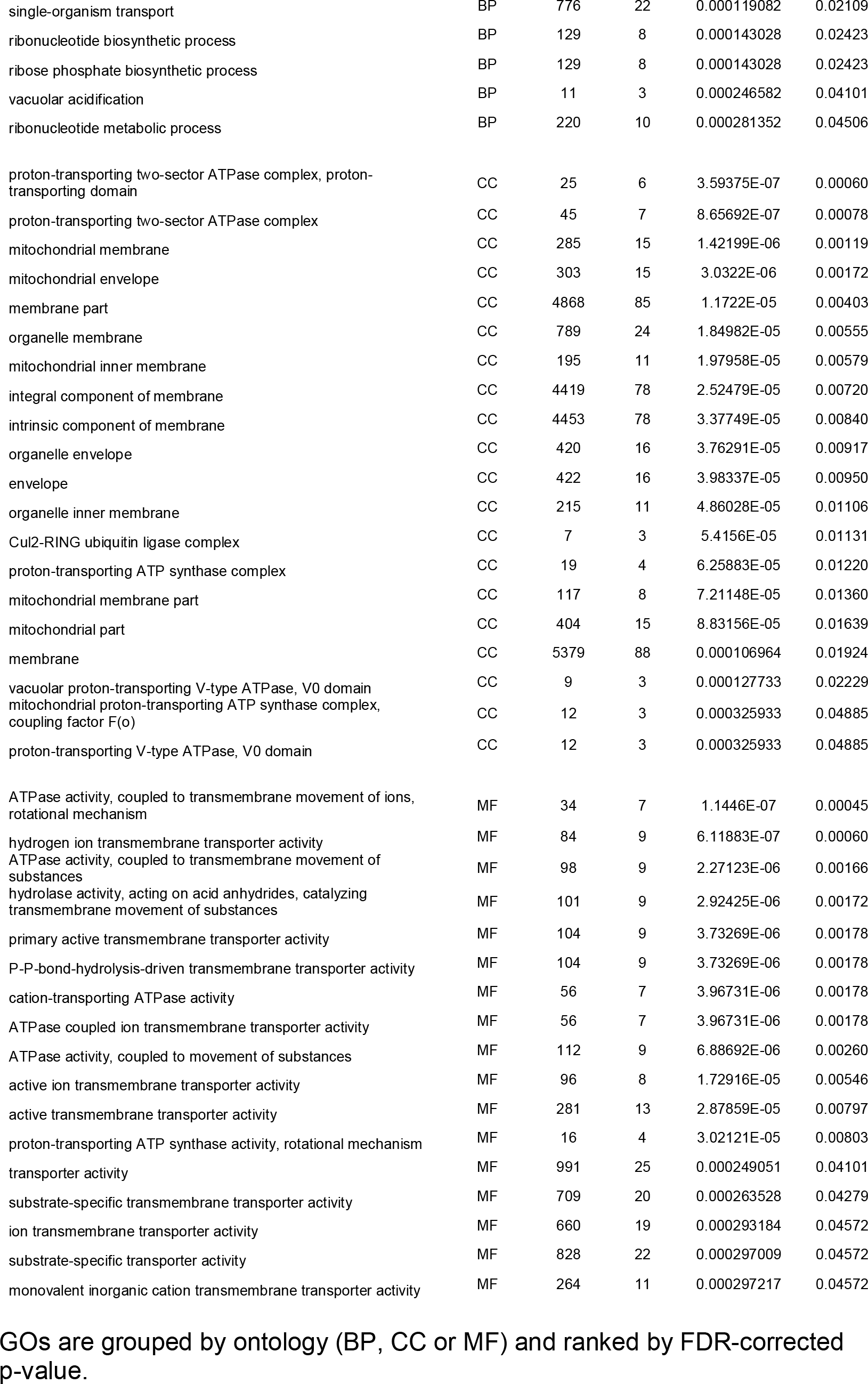

## List of Abbreviations

AD: Alzheimer’s disease
ATP: adenosine triphosphate
BP: biological process (GO term)
CC: cellular component (GO term)
DE genes: differentially expressed genes
EOfAD: early onset familial Alzheimer’s disease
FDR: false discovery rate
GEO: Gene Expression Omnibus
GO: gene ontology
MAPT: MICROTUBULE-ASSOCIATED PROTEIN TAU (human protein)
MF: molecular function (GO term)
mg: milligrams
MRI: magnetic resonance imaging
*PSEN1*: *PRESENILIN 1* (human gene)
PSEN1: PRESENILIN 1 (human protein)
*psen1*: *presenilin 1* (zebrafish gene)
Psen1: Presenilin 1 (zebrafish protein)

## Declarations

### Ethics approval and consent to participate

This study was conducted under the auspices of the Animal Ethics Committee of the University of Adelaide, under permits S-2014-108 and S-2017-073.

### Consent for publication

Not applicable

### Availability of data and material

The datasets generated and/or analysed during the current study are available in the GEO repository (https://www.ncbi.nlm.nih.gov/geo/) under accession number GSE126096.

*psen1*^*Q96_K97del*^ mutant zebrafish are available upon request. However, due to Australia’s strict quarantine and export regulations, export of fish involves considerable effort and expense and these costs must be borne by the party requesting the fish.

## Authors’ contributions

MN conceived the project, sought funding, generated the *psen1*^*Q96_K97del*^ mutant zebrafish, identified the genotype of individuals, and isolated mRNA from zebrafish brains. NH processed the RNA-seq data and performed bioinformatics analysis to identify DE genes and GOs. SP supervised the work of NH and performed data quality checks. ML conceived the project, sought funding, coordinated the project, and drafted this research report. All authors contributed to interpretation of data and to reviewing and editing drafts of the submitted manuscript.

## Acknowledgements

The authors wish to thank the Carthew Family Foundation and Prof. David Adelson for their encouragement and support.

## Tables and additional files

### Information regarding <u>Additional File 1</u>

This is a Microsoft Excel spreadsheet file with the name, “Additional File 1.xlsx”.

### Title of data

Genes differentially expressed between heterozygous mutant and wild type brains at 6 months

### Description of data

Additional File 1 lists the genes identified as differentially expressed between the brains of heterozygous *psen1*^*Q96_K97del*^ mutant fish and the brains of their wild type siblings at an age of 6 months. Genes are ranked according to FDR-corrected p-value. Only genes with a FDR-corrected p-value less than 0.05 are shown. “FC” denotes fold change. “DE” denotes differential expression. For DE_Direction, “1” denotes increased expression in the mutant and “-1” denotes decreased expression in the mutant.

